# Tonic GluD1 channel current is independent of G protein activity in the dorsal raphe nucleus

**DOI:** 10.1101/2022.01.20.477055

**Authors:** Daniel S. Copeland, Stephanie C. Gantz

## Abstract

Previously, using electrophysiological recordings from adult male and female mouse brain slices containing the dorsal raphe nucleus, we showed that GluD1_R_ channels carry ionic current and are modulated via activation of Gα_q_-coupled α1-adrenergic receptors (α1-A_R_) in a GTP-dependent manner (Gantz et al., 2020). GluD1_R_ channels also carry a tonic cation current, generally ~−20 pA at subthreshold membrane potentials (Gantz et al., 2020). The origin of tonic GluD1_R_ channel current is unknown. Here, using the same preparation, we show there is no role of on-going G protein-coupled receptor activity in generating or sustaining tonic GluD1_R_ channel current. Neither augmentation nor disruption of G protein activity had an effect on tonic GluD1_R_ current. These results reveal that tonic GluD1_R_ current arises from a mechanism separate from on-going activity of G protein-coupled receptors. Under current clamp, block of GluD1_R_ channels hyperpolarized the membrane by ~10 mV at subthreshold potentials leading to reduced excitability. Thus, GluD1_R_ channels carry a G protein-independent tonic current that contributes to subthreshold drive of action potential firing in the dorsal raphe nucleus.

## Introduction

The majority of excitatory neurotransmission in the central nervous system is produced by ionic current carried by the ionotropic glutamate receptors (iGluRs). Lesser known in the iGluR family are the delta glutamate receptors (GluD1_R_ and GluD2_R_) which share <30% amino acid sequence identity with the other family members (Araki et al., 1993; Lomeli et al., 1993). GluD1_R_ are expressed widely in the brain (Hepp et al., 2015; Konno et al., 2014; Nakamoto et al., 2020), where they regulate inhibitory and excitatory synapse formation and composition in complex with trans-synaptic and secreted proteins (Dai et al., 2021; Fossati et al., 2019; Gawande et al., 2021; Tao et al., 2018). These functions do not involve ion conduction through the channel pore (reviewed in Yuzaki and Aricescu, 2017). The study of ion channel function of GluD1_R_ has been limited since there is no known agonist that binds to GluD1_R_ directly to gate opening of the channel. Nonetheless, we and others have demonstrated that GluD1_R_ and GluD2_R_ channels carry ionic current upon activation of Gα_q_ protein-coupled receptors (GqPCRs), either metabotropic glutamate (mGlu_R_, Ady et al., 2013; Benamer et al., 2018; Dadak et al., 2017) or α1-adrenergic receptors (Gantz et al., 2020), through a process that involves intact G protein signaling. Intriguingly, in cell lines and brain slices, GluD1_R_ and GluD2_R_ channels are open in the presumed absence of agonists and carry tonic cation current (Gantz et al., 2020; Lemoine et al., 2020). The origin of the tonic GluD_R_ current is unknown.

Typically, GPCRs are activated when extracellular ligands bind to the receptor and force a conformational change which initiates downstream signal transduction mechanisms. In principle, GqPCRs could exhibit low levels of activation in response to ambient ligand, as demonstrated recently for Gα_i/o_ protein-coupled dopamine D2 receptors (Rodriguez-Contreras et al., 2021). GPCRs can also be constitutively active; entering an active state conformation in the absence of ligand (reviewed in Bond and IJzerman, 2006). Despite knowledge that GluD1_R_ channels are modulated by a GTP-dependent mechanism (Gantz et al., 2020) whether the tonic GluD1_R_ current is a product of low-level GqPCR activity is not established.

Here, using patch-clamp electrophysiology in acute mouse brain slices containing the dorsal raphe nucleus, we show that manipulating G protein activity did not impact the amplitude of the tonic GluD1_R_ current. Augmentation of on-going G protein activity amplified tonic potassium current carried by G protein-coupled inwardly rectifying potassium (GIRK) channels, but not tonic GluD1_R_ current. Inverse agonism of α1-adrenergic receptors did not affect the tonic GluD1_R_ current, suggesting a key modulator of GluD1_R_ channel current is not responsible for generating tonic GluD1_R_ current. Further, depletion of cell-autonomous G protein activity did not change the amplitude of the tonic GluD1_R_ current. Thus, tonic GluD1_R_ current arises from a mechanism separate from on-going, cell-autonomous GPCR activity.

## Results and Discussion

### GluD1_R_ channels carry a tonic current

Whole-cell voltage-clamp recordings were made from dorsal raphe neurons in acute brain slices from wild type mice at 35° C in the presence of NMDA_R_, AMPA_R_, and kainate_R_ blockers, using a potassium-based internal solution (V_hold_ −65 mV). Our previous work showed that GluD1_R_ channels carry an ~−20 pA tonic current, revealed by the application of a channel blocker, 1-naphthyl acetyl spermine (NASPM) and by genetic deletion of the GluD1_R_ channel (Gantz et al., 2020). In agreement with our prior work, here we show that application of NASPM (100 μM) produced an apparent outward current of 18.2±2.8 pA (p = 0.02, n = 7, Figures 1A and B). On average, the current peaked in 3.5 min and reversed in 11 min upon washout of NASPM. The outward current was accompanied by an increase in the membrane resistance (p = 0.02, n = 7, Figure 1C) and a reduction in the membrane noise variance (σ^2^, p = 0.02, n = 7, Figure 1D), indicating fewer open channels. NASPM failed to change the current when external Na^+^ (126 mM) was replaced with N-methyl D-glucamine (−3.0±3.6 pA, p = 0.50, n = 3). In all, these findings reproduce those of our previous work (Gantz et al., 2020) and demonstrate that NASPM blocks a tonic, sodium-dependent inward current.

**Figure 1.**
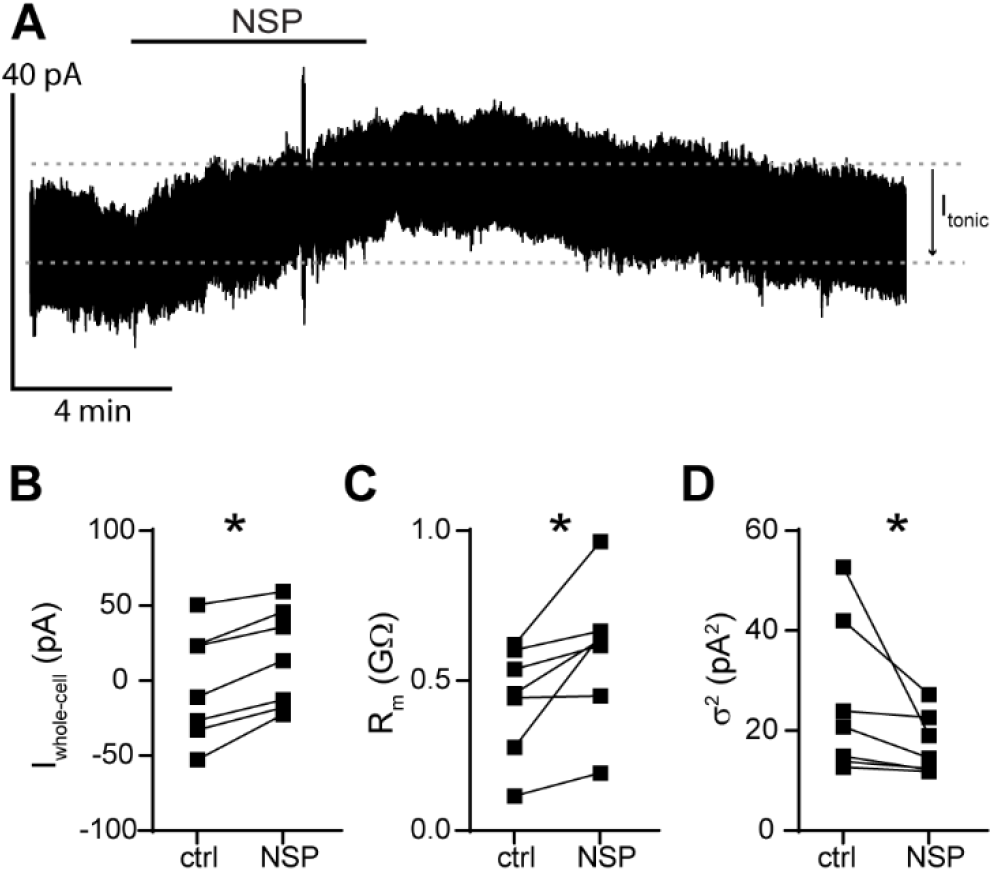
NASPM reveals a tonic inward current carried by GluD1_R_ channels. (**A**) Representative whole-cell voltage-clamp recording of the apparent outward current produced by application of NASPM (NSP, 100 μM). Dashed lines indicate baseline current (bottom) and the peak of the outward current (top). Tonic current was measured as the difference of these lines (I_tonic,_ arrow). (**B**) Plot of the whole-cell current (V_hold_ −65 mV) in control conditions (ctrl) and after application of NASPM (NSP, p = 0.016, n = 7). (**C**) Plot of basal membrane resistance recorded in control conditions (ctrl) and during NASPM application (NSP p = 0.016, n = 7). (**D**) Plot of membrane noise variance (σ^2^) in control conditions (ctrl) and during NASPM application (NSP, p = 0.016, n = 7). Line and error bars represent mean ± SEM. * denotes statistical significance.

### Tonic GluD1_R_ current is not produced by G protein activity

GluD1_R_ and GluD2_R_ channels carry ionic current following the activation of either mGlu_R_ or α1-adrenergic receptors (Ady et al., 2013; Benamer et al., 2018; Dadak et al., 2017; Gantz et al., 2020; Lemoine et al., 2020). GluD2_R_ current, seen upon mGlu_R_1 activation in cell lines, is dependent on canonical GqPCR signaling as the agonist-induced current is blocked by bath application of Gα_q/11_ or phospholipase C inhibitors (Dadak et al., 2017). Similarly, GluD1_R_ ionic current activated by α1-adrenergic receptors in dorsal raphe neurons is abolished after internal dialysis with GDPβS-Li_3_ (Gantz et al., 2020), a non-specific disruptor of G protein activity and other processes requiring a GDP-GTP exchange. Following our original report of tonic GluD1_R_ current in brain slices (Gantz et al., 2020), Lemoine et al., (2020) reported a very similar tonic current carried by GluD2_R_ channels when expressed in cell lines with mGlu_R_1. mGlu_R_, like many GPCRs, can exhibit constitutive activity in the absence of agonist (Prézeau et al., 1996) and low-level constitutive activity of GPCRs affects other subthreshold cation conductances (Lu et al., 2010; Philippart and Khaliq, 2018; Quallo et al., 2017; Shen et al., 2012; Zhang et al., 2012). But, the involvement of GqPCRs in generating tonic GluD1_R_ current has not been explored.

To test whether increased G protein activity was sufficient for generating tonic GluD1_R_ current, the internal recording solution was supplemented with a non-hydrolyzable GTP analog (guanosine-5’-[( β,γ)-imido]triphosphate, GppNHp, 1 mM), which binds irreversibly to Gα and elevates G protein activity. In dorsal raphe neurons, dialysis with GppNHp produces a tonic outward current carried by G protein-coupled inwardly rectifying potassium (GIRK) channels (Loucif et al., 2006) by elevating free Gβγ subunits which gate GIRK channels (Pfaffinger et al., 1985). In agreement, whole-cell dialysis of GppNHp-containing internal solution (≥10 min) produced a tonic outward current with a reversal potential of −108 mV (Figures 2A and B), consistent with the expected reversal potential of potassium (calculated E_K_: −104 mV). Application of BaCl_2_ (100 μM), which blocks GIRK channels (Gantz et al., 2013) produced an apparent inward current (−46.0±10.6 pA, Figure 2A) with GppNHp-but not GTP-containing internal solution (p = 0.0003, n = 9-16, Figure 2C). These data demonstrate that amplifying G protein signaling with GppNHp produces standing currents carried by G protein-gated ion channels, consistent with previous studies (Kramer and Williams, 2016; Loucif et al., 2006). In the continued presence of BaCl_2_, tonic GluD1_R_ current was measured following application of NASPM (Figure 2D). On average, tonic GluD1_R_ current was −16.7±2.8 pA, which was not different from current measured with GTP-containing internal solution (p = 0.72, n = 5-8, Figure 2E). BaCl_2_ had no effect on the amplitude of the α1-adrenergic receptor-dependent excitatory postsynaptic current (ctrl: −20.6±3.6 pA; BaCl_2_: −26.0±4.1 pA, p = 0.20, n = 8), evoked by electrical stimulation of the brain slice (5 pulses, 0.5 ms, 60 Hz) delivered via a monopolar stimulating electrode, indicating that at this concentration, external BaCl_2_ does not affect conductance of GluD1_R_ channels. Taken together, the data suggest that augmenting G protein activity has a negligible impact on the amplitude of the tonic current carried by GluD1_R_ channels.

**Figure 2.**
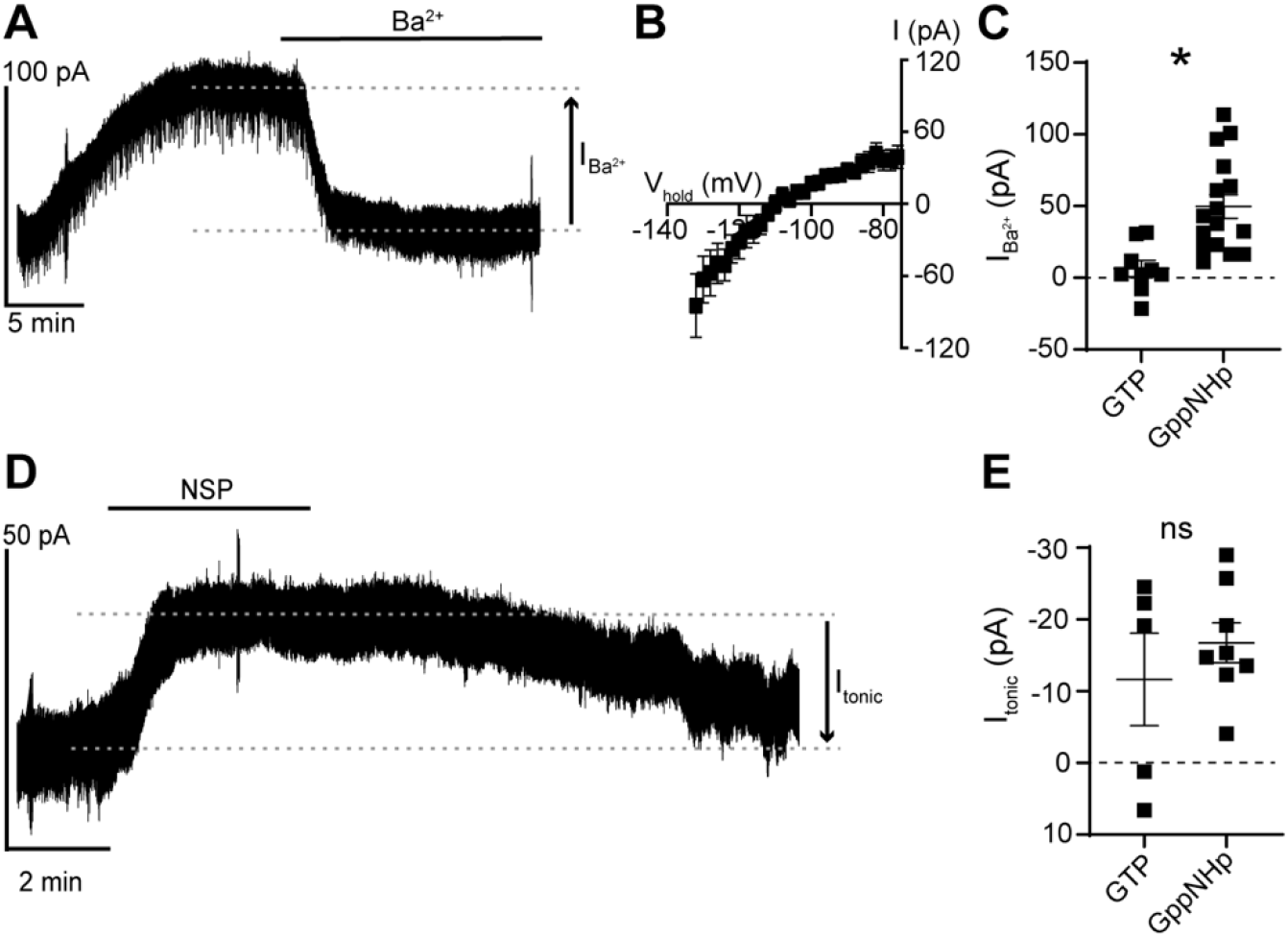
Augmentation of G protein activity with GppNHp has no effect on the tonic current carried by GluD1_R_. (**A**) GppNHp-containing internal solution produced a tonic Ba^2+^-sensitive (100 μM) outward current, shown in a representative whole-cell voltage-clamp recording. (**B**) Current-voltage relationship of the tonic Ba^2+^-sensitive (100 μM) outward current demonstrating reversal near expected E_K_ and inward rectification. (**C**) Plot of tonic GIRK currents measured in control conditions (GTP) and with GppNHp-containing internal solution (p = 0.0003, n = 9 and 16 respectively). (**D**) GppNHp had no effect on the NASPM (NSP, 100 μM)-sensitive inward current, shown in a representative recording. (**E**) Plot of the magnitude of GluD1_R_ tonic current measured with GTP-containing internal solution as compared to GppNHp-containing internal solution (p = 0.72, n = 5 and 8 respectively) when measured in external Ba^2+^ to block tonic GIRK current. Line and error bars represent mean ± SEM. * denotes statistical significance, ns denotes not significant.

In principle, GPCRs could be activated by ambient ligand in brain slices to produce a small tonic current. In midbrain dopamine neurons, ambient activation of Gα_i/o_-coupled dopamine D2 receptors produces a tonic GIRK current of ~9 pA (Rodriguez-Contreras et al., 2021). Next, we tested whether tonic GluD1_R_ current was dependent on α1-adrenergic receptor activity, either from ambient ligand or constitutive activity, by applying an α1-adrenergic receptor inverse agonist prazosin (100 nM, Hein et al., 2001). Prazosin had no effect on the magnitude of the tonic GluD1_R_ current (−21.2±2.8 pA, p = 0.22, n = 5). In midbrain dopamine neurons, GluD1_R_ channel current is produced by activation of mGlu_R_ (Benamer et al., 2018), suggesting a similar mechanism may occur in the dorsal raphe. Moreover, in the dorsal raphe, Gα_q_-coupled histamine H_1_ and orexin OX_2_ receptors converge on the same downstream effectors as α1-adrenergic receptors (Brown et al., 2002), suggesting if the tonic GluD1_R_ current was produced via G protein activity, there are many types of receptors to consider. As a broad test as to whether tonic GluD1_R_ current was dependent on G protein signaling, recordings were made with an internal solution where GTP was replaced with a non-hydrolyzable analog of GDP, GDPβS-Li_3_ (1.24 mM) which acts as a competitive antagonist at GTP-binding sites. Within 10 mins of whole-cell dialysis with GDPβS-containing internal solution, application of noradrenaline (30 μM) produced an inward GluD1_R_ current (Figures 3A and B). By ≥20 min of whole-cell dialysis, the noradrenaline-induced current was abolished (p = 0.02, n = 7, Figures 3A and B), confirming efficacy of GDPβS to arrest α1-adrenergic receptor-GluD1_R_ channel signaling (Gantz et al., 2020). In contrast, GDPβS had no effect on the tonic GluD1_R_ current when compared with GTP-containing internal solution with or without a similar concentration of LiCl (GDPβS-Li_3_: −28.7±4.9 pA, n = 13, GTP: −19.3±2.7 pA, GTP + LiCl: −23.5±3.9 pA, n = 8-13, p = 0.46, Figure 3C). Further, there was no decrement in the magnitude of the tonic GluD1_R_ current with repeat applications of NASPM (p = 0.84, n = 6, Figures 3D and E). Taken together, these results demonstrate that tonic GluD1_R_ current is independent of cell-autonomous G protein signaling.

**Figure 3.**
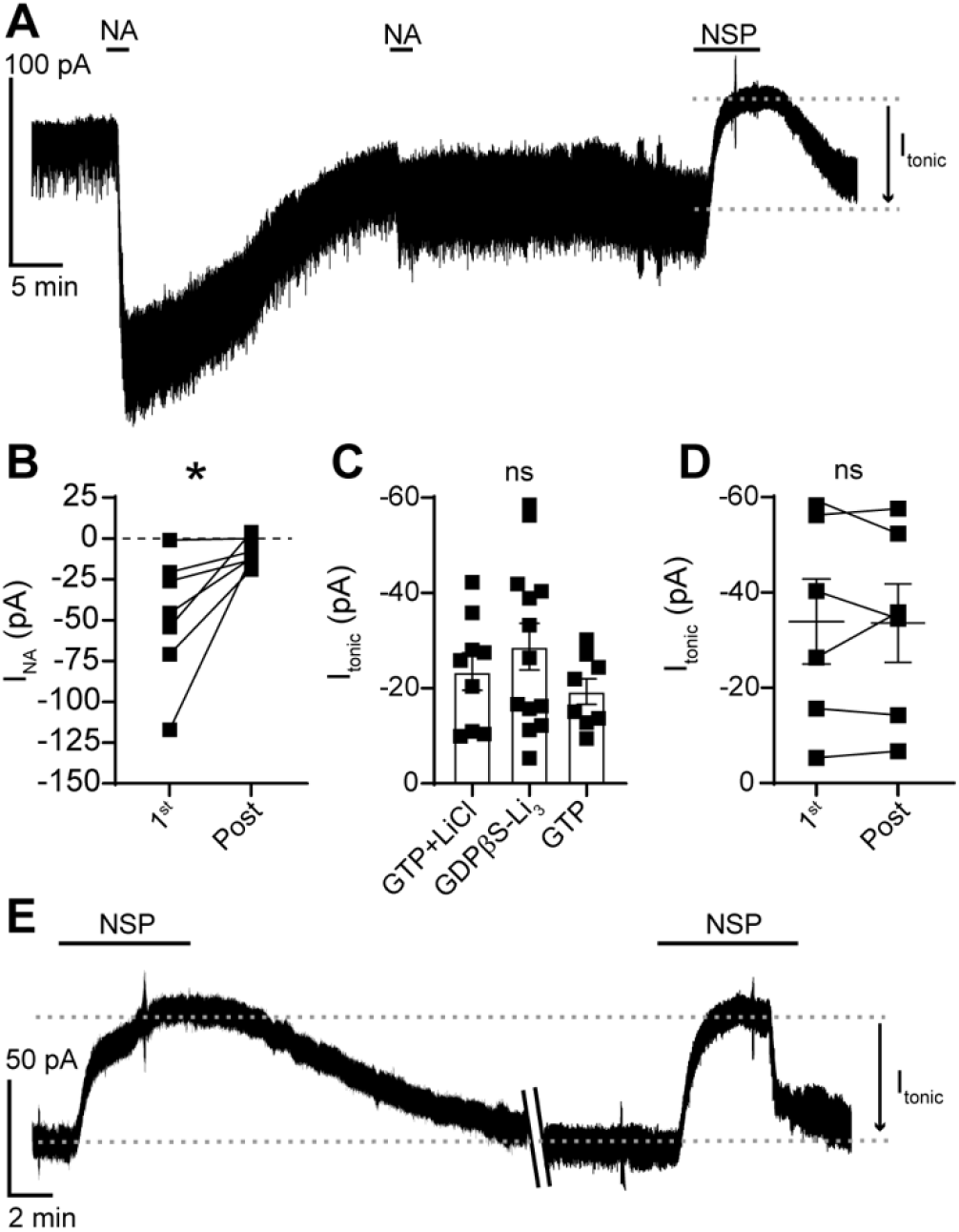
The GluD1_R_ tonic current is not dependent on G protein signaling. (**A**) With GDPβS-containing internal solution, noradrenaline-induced GluD1_R_ current (I_NA_) was diminished by >20 minutes post-dialysis, shown in a representative trace. (**B**) With GDPβS-containing internal solution, the amplitude of I_NA_ ran down with whole-cell dialysis; shown in a plot of the first application of noradrenaline (30 μM, 1^st^) compared to application of noradrenaline >20 min post-dialysis (Post, p = 0.02, n = 7). (**C**) Plot of I_tonic_ measured after dialysis with GTP + LiCl−, GDPβS-Li_3−_, and GTP-containing internal solution, displaying no significant difference between the groups (p = 0.46, n = 9, 13, 8 respectively). (**D**) Plot of I_tonic_ recorded with GDPβS-containing internal solution for the first application of NASPM (1^st^) and application of NASPM >45 min post-dialysis, showing no difference in the average amplitude (Post, p = 0.84, n = 6). (**E**) With GDPβS-containing internal solution, repeated application of NASPM revealed tonic GluD1_R_ current without a decrement in amplitude, shown in a representative trace. Slanted lines indicate a 40-minute wash time in the recording. Line and error bars represent mean ± SEM. * denotes statistical significance, ns denotes not significant.

### Tonic GluD1_R_ current provides subthreshold drive of action potential firing

Throughout the central nervous system, many types of neurons fire action potentials in a rhythmic ‘pacemaker’ pattern. Some are autonomous pacemakers, driven by intrinsic membrane properties, while others are conditional pacemakers that rely on synaptic input and receptor stimulation. A common feature in autonomous pacemakers is the presence of a tonic, subthreshold, tetrodotoxin-insensitive, cation/sodium current (Eggermann et al., 2011; Jackson et al., 2004; Khaliq and Bean, 2010; Li et al., 2021; Lu et al., 2007; Raman et al., 2000). While many different types of channels are involved, these tonic currents each function to depolarize the membrane to ~−60 mV where voltage-dependent mechanisms of action potential firing are engaged. Primarily, serotonin neurons are conditional pacemakers and require subthreshold drive from noradrenergic afferents and activation of α1-adrenergic receptors (Baraban et al., 1978; Vandermaelen and Aghajanian, 1983) much like other conditional pacemakers which require activation of Gα_q_-coupled orexin or muscarine receptors (Egorov et al., 2019, 2002; Top et al., 2004; Yamada-Hanff and Bean, 2013). In these neurons, activation of GqPCRs leads to subthreshold (~−70 to −55 mV) depolarization via a very similar cation current as the tonic current observed in autonomous pacemakers. While these tonic cation currents are essential for subthreshold depolarization, it is not unusual for the current to be quite small, only a few to tens of picoamperes (Jackson et al., 2004; Raman et al., 2000; Taddese and Bean, 2002).

To determine if tonic GluD1_R_ current contributed to subthreshold excitation, whole-cell current clamp recordings were made from dorsal raphe neurons and APs were evoked with somatic current injection (1.5 s, 20 pA increments, Figure 4A). In the absence of noradrenaline, dorsal raphe neurons are silent or fire slowly and erratically (Baraban et al., 1978; Svensson et al., 1975). Consistent with the absence of ambient noradrenaline in brain slices, 5/10 neurons were firing spontaneously at a slow and irregular rate (0.8±0.3 Hz) and 5/10 only fired in response to current injection. After application of NASPM (30 – 50 μM), 1/10 neurons fired spontaneously, and the rest became quiescent until APs were evoked by current injection (Figure 4B). At subthreshold potentials (−80 to −55 mV), NASPM hyperpolarized the membrane by ~10 mV (Figure 4C), consistent with expectations given the magnitude of the tonic GluD1_R_ current (−19.6±4.0 pA) and basal membrane resistance (555.6±45.4 MΩs).

**Figure 4.**
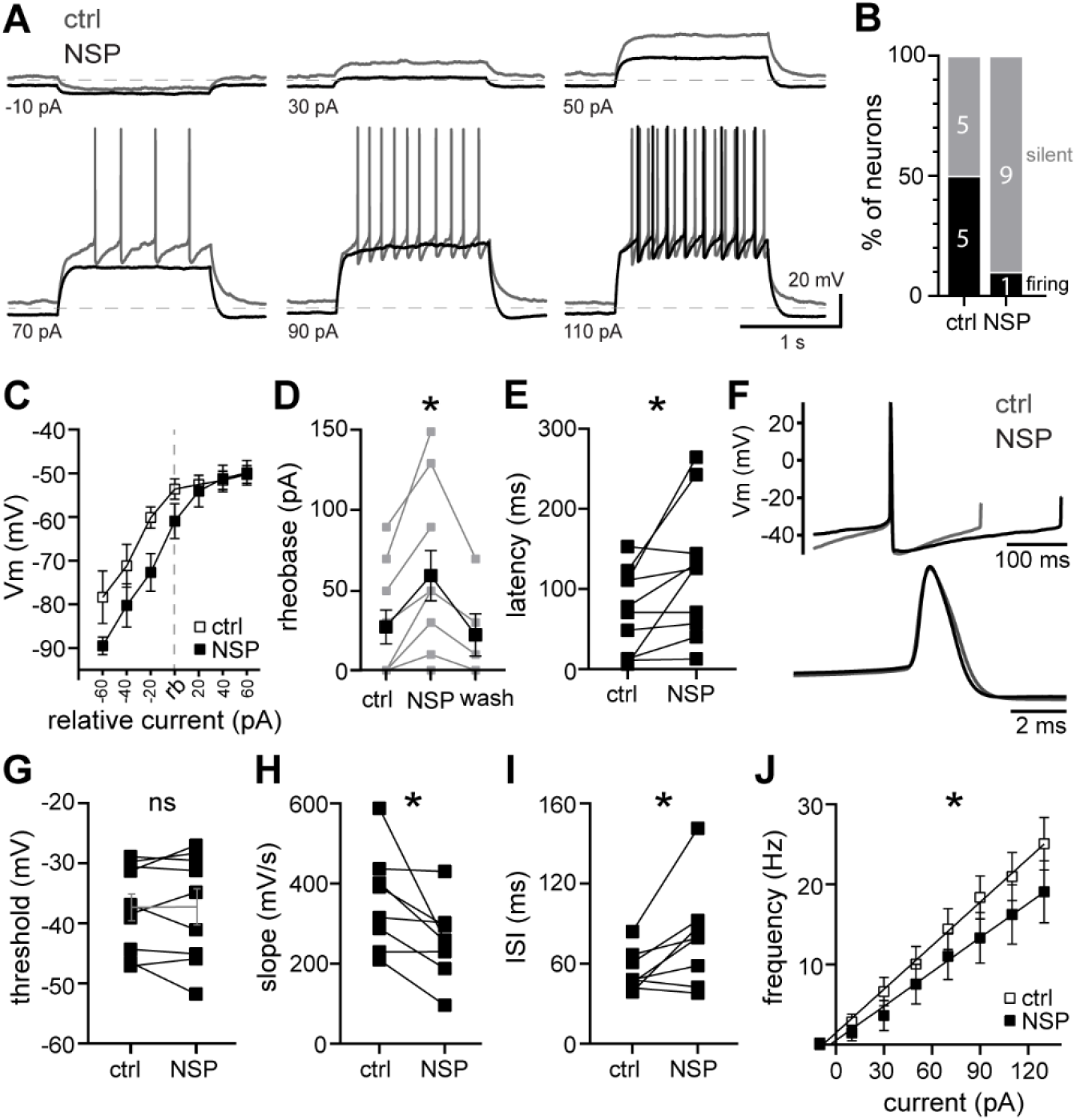
Tonic GluD1_R_ current provides subthreshold drive of action potential firing. (**A**) Representative traces of whole-cell current clamp recordings of membrane potential and AP firing evoked by current injection (1.5 s) demonstrating hyperpolarization by NASPM. Dashed line is at −80 mV. (**B**) Distribution of firing response in control (ctrl) and after application of NASPM (NSP). In control conditions 5/10 neurons fired spontaneously without current injection and 5/10 were silent. In the same neurons after NASPM, 1/10 fired spontaneously and 9/10 were silent. (**C**) Plot of the membrane potential (Vm) versus injected current relative to rheobase (rb) in control (open squares) and NASPM (closed squares), demonstrating that NASPM produced a hyperpolarization at subthreshold potentials. Membrane potential during AP firing was measured as the average interspike Vm. (**D**) Plot of the minimum current needed to induce firing (approximate rheobase) in control conditions, NASPM, and following wash out of NASPM (10 min) (p = 0.005, n = 5-10). (**E**) NASPM increased the latency to fire the first AP upon current injection (150 pA, p = 0.01, n = 10). (**F**) Average AP waveform recorded in control and in NASPM, aligned at peaks. Below, expanded timescale. (**G**) NASPM had no effect on the AP threshold (150 pA, measured from the 2^nd^ AP, p < 0.99, n = 9). (**H**) NASPM decreased the slope of the voltage trajectory between APs (90 pA, measured in the middle 60% of the interspike interval of the first 5 APs, p = 0.02, n = 8). (**I**) NASPM increased the interspike interval during evoked firing (90 pA, averaged from first 5 APs, p = 0.04, n = 8). (**J**) Plot of the initial firing frequency (first 3 APs) versus injected current in control (open squares) and NASPM (closed squares). Line and error bars represent mean ± SEM. * denotes statistical significance, ns denotes not significant.

Consequently, NASPM increased the minimum current necessary to evoke AP firing (approximate rheobase), which reversed upon 10 min washout of NASPM (p = 0.005, n = 4-9, Figure 4D) and increased the latency to fire the first AP (p = 0.02, Figure 4E). In contrast, once the membrane reached threshold, NASPM had little-to-no effect on average membrane potential between APs (Figure 4C). The AP waveform was largely unaffected by NASPM (Figure 4F), consistent with a reversal potential of ~−30 mV and intrinsic inward rectification (Gantz et al., 2020).

However, it should be noted that NASPM preferentially blocks inward flow and strong depolarization relieves pore block (Koike et al., 1997). Thus, observation of any contribution of outward ion flux may be obscured. There were no differences in the AP half-width (p=0.19), after-hyperpolarization (p = 0.38), height (p = 0.06), or threshold (p > 0.99, Figure 4G). But there was a significant decrease in the slope of the voltage trajectory between APs (p = 0.02, Figures 4F and H), resulting in a delay to the next AP (interspike interval, p = 0.04, Figure 4I). Overall, NASPM reduced AP firing frequency (p = 0.01, two-way ANOVA, measured from the first 3 APs, Figure 4J). Thus, tonic GluD1_R_ current contributes to subthreshold drive of action potential firing. Provided the widespread distribution in the brain (Hepp et al., 2015; Konno et al., 2014; Nakamoto et al., 2020), GluD1_R_ channels may contribute to pacemaking in other neuronal populations, whether via intrinsic tonic current or following GqPCR activation. Future work will be needed to determine whether tonic GluD1_R_ current arises from an intrinsic property of the channel (i.e. an open *apo*-state) or is generated by an unidentified ligand.

## Methods

### Animals

All studies were conducted in accordance with the University of Iowa with the approval of the University of Iowa Institutional Animal Care and Use Committee. Male and female wild type C57BL/6J (>2 months old, The Jackson Laboratory, #000664) mice were used. Mice were group-housed on a 12:12 h light cycle.

### Brain slice preparation and electrophysiological recordings

Brain slices and electrophysiological recordings were made as previously described (Khamma et al., 2021). In brief, mice were deeply anesthetized with isoflurane and euthanized by decapitation. Brains were removed and placed in warmed and bubbled (95/5% O_2_/CO_2_) modified Krebs’ buffer containing (in mM): 126 NaCl, 2.5 KCl, 1.2 MgCl_2_, 1.2 CaCl_2_, 1.2 NaH_2_PO_4_, 21.5 NaHCO_3_, and 11 D-glucose with 5 μM MK-801 to reduce excitotoxicity and increase slice viability. In the same solution, coronal dorsal raphe slices (240 μm) were obtained using a vibrating microtome (Leica 1220S) and incubated at 28 °C >30 minutes prior to recording.

Electrophysiological recordings were made in NBQX (3 μM) solutions at 35 °C with Multiclamp 700B amplifiers (Molecular Devices), Digidata 1440A and 1550B A/D converters (Molecular Devices), and Clampex software (Molecular Devices) with borosilicate glass electrodes (World Precision Instruments) wrapped with Parafilm to reduce pipette capacitance. Pipette resistances were 3.8 to 4.5 MΩ when filled with an internal solution containing, (in mM) 104.56 K-methylsulfate, 3.73 KCl, 5.3 NaCl, 4.06 MgCl_2_, 4.06 CaCl_2_, 7.07 HEPES (K), 3.25 BAPTA (K4), 0.26 GTP (sodium salt), 4.87 ATP (sodium salt), 4.59 creatine phosphate (sodium salt), pH 7.24 with KOH, mOsm ~274, for whole-cell patch-clamp recordings. Series resistance was monitored throughout the experiment. Reported voltages are corrected for a liquid junction potential of −8 mV between the internal and external solution. All drugs were applied via the patch-pipette or by bath application. Noradrenaline was applied in the presence of an α2-adrenergic antagonist, idazoxan (1 μM).

### Materials

BaCl_2_: Sigma, 217565

GDPβS-Li_3_: Sigma, G7637

GppNHp: Sigma, G0635

Idazoxan: Sigma, I6138

LiCl: Sigma, L4408

MK801: Tocris, 0924

NASPM: Tocris, 2766

NBQX: Tocris, 1044

NMDG: Sigma, M2004

Noradrenaline: Tocris, 5169

Prazosin: Tocris, 623

Igor-Pro 6.37: Wavemetrics

DataAccess: Bruxton Corporation

Clampfit 10.7: Molecular Devices

GraphPad Prism 8.4.3: GraphPad Software

### Experimental Design and Statistical Analysis

Data were analyzed using Clampfit 10.7 or Igor-Pro 6.37 software, and are presented in representative traces, scatter plots and bar graphs with means ± SEM. Unless otherwise noted, *n* = number of cells as biological replicates. Tonic current was measured as the peak of the NASPM-induced current minus the whole-cell current prior to NASPM application. Reversal potentials were determined using linear fit of the averaged data accounting for scatter amongst the replicates using five data points above and below where the current reversed polarity. Significant differences were determined via Wilcoxon matched-pairs signed rank tests for within-group comparisons, and Mann Whitney tests for between-group comparisons. A difference of p<0.05 was considered significant. Exact values are reported unless p<0.0001 or >0.999. Statistical analysis was performed using GraphPad Prism 8.4 (GraphPad Software, Inc.).

## Acknowledgements

This research was funded by a startup award from the University of Iowa Carver College of Medicine to S.C.G. We thank Holly S. Hake for critical reading and comments on the manuscript.

## Author Contributions

Conceptualization, S.C.G., experimental setup and development, D.S.C. and S.C.G.; experiments, D.S.C. and S.C.G.; analysis, D.S.C. and S.C.G.; writing & editing, D.S.C. and S.C.G.

